# A Scalable and Adaptive Ultra-high-density Fan-out Strategy for High-throughput Flexible Microelectrodes

**DOI:** 10.1101/2022.11.07.515530

**Authors:** Huiling Zhang, Yang Wang, Xinze Yang, Miao Yuan, Xiaowei Yang, Qiang Gui, Yijun Wang, Hongda Chen, Ruping Liu, Weihua Pei

**Author notes:** Huiling Zhang and Yang Wang contributed equally to this work.

## Abstract

Flexible neural microelectrodes demonstrate higher compliance and better biocompatibility than rigid electrodes. They have multiple microfilaments can be freely distributed across different brain regions. However, high-density fan-out of high-throughput flexible microelectrodes remains a challenge since monolithic integration between electrodes and circuits is not currently available as it is for silicon electrodes. Here, we proposed a high-density fan-out strategy for high-throughput flexible electrodes. The flexible electrodes are partially overlapped on the printed circuit board (PCB). A modified wire bonding method is used to reduce the pads area on the PCB. Both the traces and pads on the PCB are optimized to minimize the back-end package. It is significantly reducing the connection area. In addition, the vertical rather than horizontal connection between the connector and the PCB further decrease the volume of the package. 1024 channel flexible electrodes are bonded to the high-density PCB within an area of 11.8×10mm^2^, and the success connection rate can reach 100%. The high-density fan-out strategy proposed in this paper can effectively reduce the volume of high-throughput flexible electrodes after packaging, which facilitates the miniaturization of in vivo multichannel recording devices and contributes to long-term recording.

## 1 Introduction

Flexible neural microelectrodes are an essential tool for large-scale and long-term neural recordings. Low Young’s modulus allows flexible electrodes to be better matched to soft brain tissue. Compared with the rigid silicon-based electrode, flexible electrodes dramatically reduce chronic immune response after implantation, allowing for long-term, high-quality recordings [1–7]. High-throughput electrodes have hundreds or even thousands of recording sites, which can capture more neuronal activity. They play an important role in clarifying the correspondence between neural activity and behavior [8–12]. High-throughput electrodes usually have thousands of sites distribute in multiple shanks, which allow simultaneous recording in adjacent brain regions. The high-throughput flexible electrodes, based on their excellent flexibility, can be freely distributed among various target brain regions according to users’ need, which shows a wide range of application prospects [13].

However, the electrophysiological signals recorded extracellularly are usually weak and need to be amplified before subsequent digital processing [14,15]. Therefore, the electrode must have a physically connection with the amplifier first. There are two types of connections, direct and indirect, as shown in Fig.1A. The direct interconnection integrates electrodes and amplifiers through a CMOS process to achieve a high-density fan-out [16,17], but this method is currently not applicable to flexible electrodes. The indirect one is usually based on a welding process that first connects the electrodes to the chip or connector and then to the data acquisition system. The amplifiers are integrated in the chip or system. The interconnection of the electrode to the chip/connector determines the final volume of the back-end package. However, the electrode sites and the pins of chip/connector hardly can be connected directly due to the mismatch in size and arrangement. APCB is often designed as a bridge between the electrodes and the chip/connector. A typical PCB can be divided into three parts (Fig.1B): the area for bonding electrodes (Area-1), the area for welding chip/connector (Area-2), and the area for metal traces and electronic components (Area-3).

**Fig.1.**
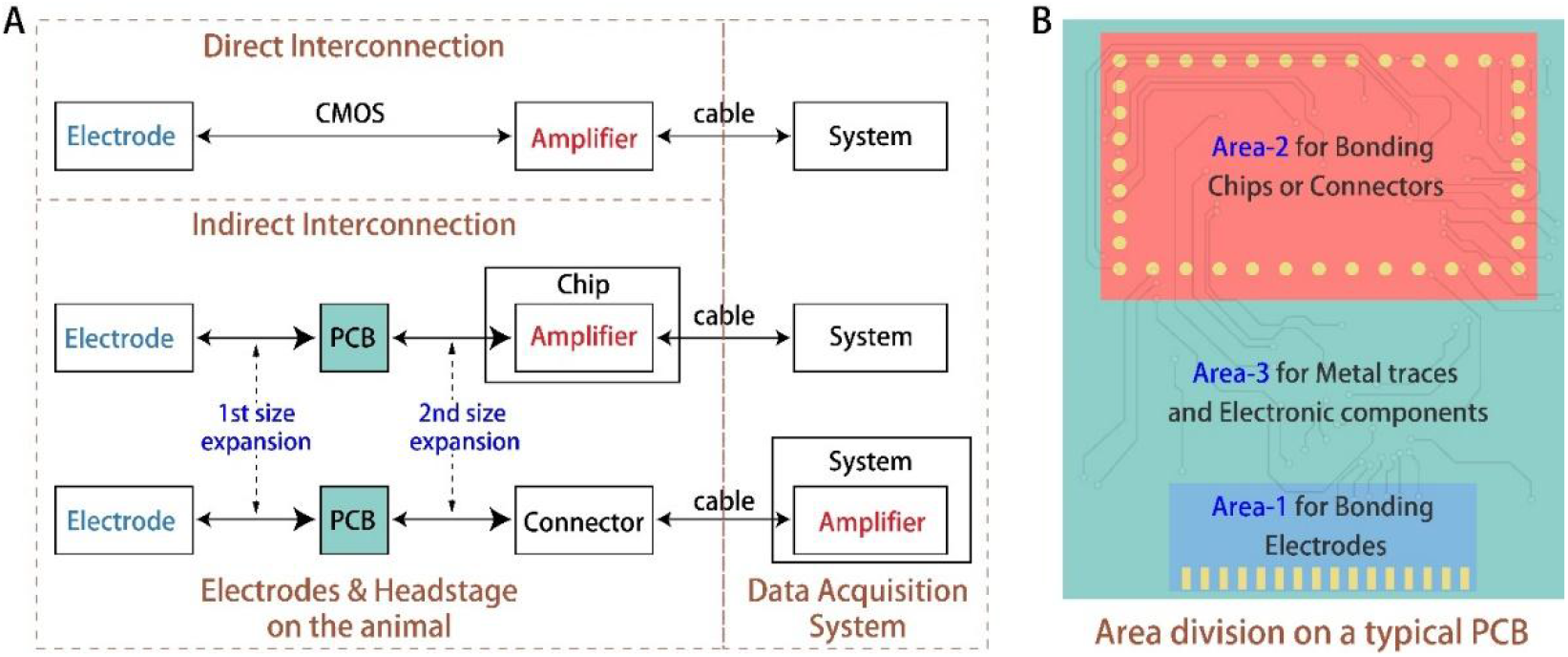
Neural microelectrodes packaging. (A) Two types of electrode connections. (B) Ares division on a typical PCB.

In Area-1, the electrode’s connection is usually realized by wire bonding, anisotropic conductive film, or flip-chip bonding. Limited by these bonding methods, the electrode pads must reach a certain size to form reliable electrical connections on the PCB. Therefore, a size expansion will occur between the back-end pads and the front-end sites on the electrode (Fig.1). In Area-2, the size and spacing of the connector pins are usually larger than the electrode pads. Therefore, there will be another size expansion between Area-1 and Area-2 on the PCB, also including the influence of Area-3 (Fig.1). This means that the size of the electrode is expanded twice during the whole fan-out packaging process, and the size expansions will aggravate nonlinearly as the electrode channels increase, due to that the metal traces from the electrode sites to the chip/connector cannot be crossed. The result is that the traditional electrode packaging is typically characterized by a low fan-out density, which can be defined as the ratio of the number of connected channels to the PCB area. For example, Pimenta et al. [18] prepared a 32-channel double-layer flexible electrode. The width of the electrode shank is only 150μm, but the width of the pad area reaches 9 mm. And after welding the connector, the back end of the whole device reaches about 15×15mm^2^. Yang et al. [19] used wire bonding to connect the PCB with a 256-channel electrode. The volume of the whole back-end package is nearly 50mm×20mm×5mm. Scholvin et al. [20] used electron beam exposure to prepare 1000-channel neural electrodes on 5 silicon needles. Although the volume of the implanted tip is about 1.5mm×50μm×5, the pad area of the electrode reaches about 30mm×17mm. When all 1000 channels are bonded to the PCB, the size of the back-end package is enlarged to 93mm×45mm, thousands of times larger than that of the implanted part. To reduce the back-end volume and increase the flexibility of the application for freely moving animals, Roso et al. [11] stacked four 256-channel electrodes together to achieve a 1024-electrode array. The volume of the implanted part is only 0.75mm×1.05mm×0.756mm, but after connecting to the PCB, the electrode volume exceeds 30mm×24mm×16mm.

The essence of low fan-out density is that the effective interconnection part is too small, compared with the whole area of the PCB. To solve this problem, a reasonable PCB design is required to cope with the size expansions. However, in the traditional electrode packaging. PCBs usually need to adapt to the size and layout of the electrode pads and the chip/connector, resulting in an unreasonable layout on it. The area of the PCB has not been used effectively. In this paper, to realize an ultra-high-density fan-out strategy for high-throughput flexible microelectrodes, a 11.8×10mm^2^ PCB with high utilization rate is designed. The proportion of the effective interconnection area on the PCB is nearly 100%. The final packaged 1024-channel electrode has a volume of no more than 12 × 11 × 8 mm^3^.

## 2 Materials and Methods

### 2.1 Methods of the high-density fan-out strategy

As shown in Fig.2. An ultra-compact PCB is designed with a high-density distribution of pads on both sides (Fig.2A). All pads run through the PCB, the electrode bonding pads are on the front side and the chip/connector welding pads on the backside (Fig.2B). On the front side of the PCB, a vertical bonding method is used to connect the flexible electrodes (Fig.2A). Modified gold-ball bonding is employed to decrease the bonding area by fabricating a hole in the center of each electrode pad [21]. Each row of pads on the PCB corresponds to one flexible microfilament electrode. (Fig.2C). The bonding parts of the flexible electrodes are partially overlapped, benefit from their ultra-thinness and flexibility. The overlap placement of the flexible filaments furtherly decreases the bonding area, increase the utilization rate. All flexible microfilaments are able to sequentially bonded to the PCB, regardless of the increase of the number of electrodes. The vertical bonding method combined with overlap placement dramatically decreases the effect of the first size expansion, especially, eliminates the nonlinear growth with increasing number of channels. On the back side of the PCB, the corresponding chip or connector can be designed to match the welding pad array.

**Fig.2.**
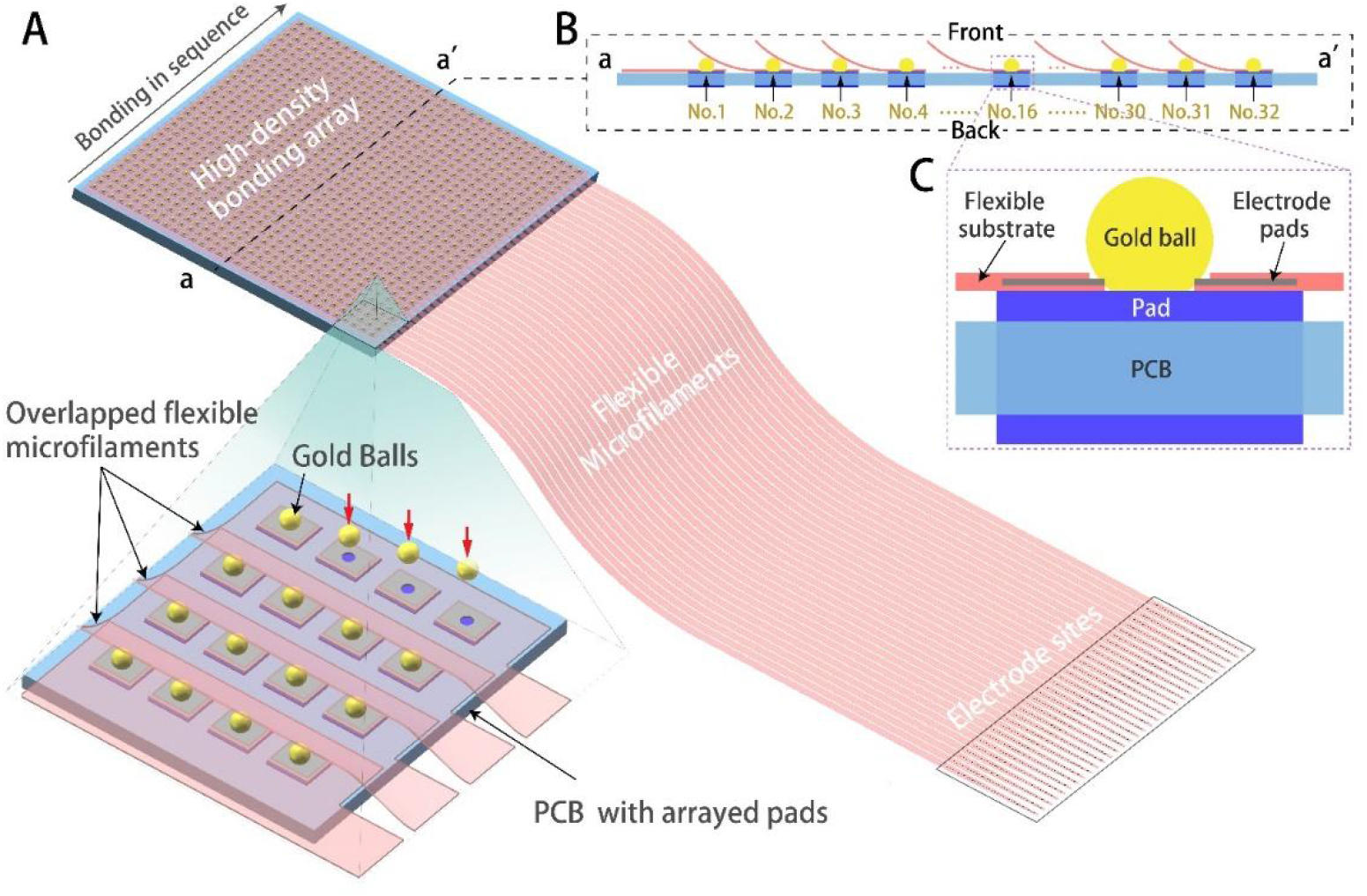
Design of the high-density fan-out strategy. (A) High-density bonding array. (B) Side of high-density bonding array. (C) Modified gold-ball bonding.

The proposed high-density fan-out method for high-throughput flexible microelectrodes can be easily scaled up because the overall package size increases linearly with the number of electrode channels.

### 2.2 Design of the high-density PCB

To validate this proposed fan-out method, we designed a PCB as shown in Fig.2A. There are 1024 pads on each side. Each bonding pad on the front side is directly connected to the welding pad on the back side face-to-face through a via hole. According to the footprints and pitch size of Omneitics connector(A79022), as well as the size of the electrode pad, each pad unit on the PCB is designed to be 280×220μm. Considering that the spacing and width of the Omnetics connector pins are 200μm, the horizontal spacing of each unit is designed to be 380μm, and the vertical spacing is 320μm to avoid short circuits between adjacent units during welding. The size of the entire pad array is 11.78×9.92mm^2^, which means that 1024 channel electrodes can be packaged in an area close to 1cm^2^.

### 2.3 The selective fan-out method

To realize the welding of the connector, eight connectors are arranged in parallel to form a 256-channel connector module (Fig.3A). It can ensure the pin spacing of all rows matches the PCB pad column spacing. The size of the entire connector module is 13×13×7mm^3^, which is close to the PCB pad array size.

**Fig. 3.**
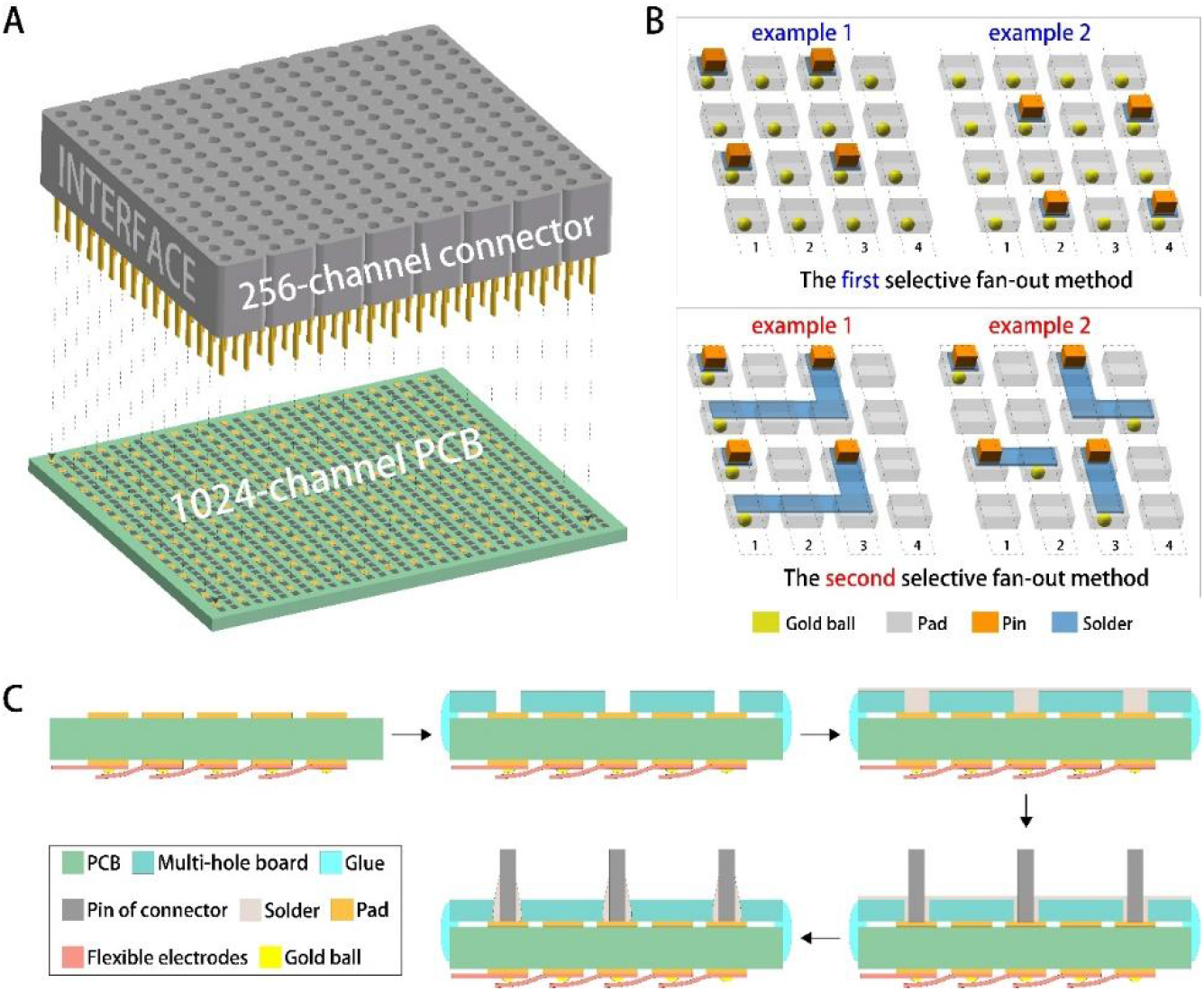
Design of the selective fan-out method. (A)256-channel connector module. (B) Two selectively fan out methods. (C) The welding process of PCB and 256-channel connector module.

Two methods were designed to selectively fan out 256 of the 1024-channel electrodes (Fig.3B). The first method is to bond all 1024 electrode pads to the bonding pad of the PCB, and by changing the position of the connector modules, 256 channels (16 flexible microfilaments with 16 channels each) can be evenly selectively connected to the connector module. The second method maintains the relative positions of the connector module and the PCB, and can achieve a more flexible interconnection by selectively bonding the electrodes to the bonding pad and shorting the welding pads of the PCB. Based on this method, it is possible to assign all the pins of a connector to one flexible microfilament, or to assign them on up to four flexible microfilaments.

### 2.4 Design of the flexible electrode

The flexible electrode needs to be adapted to the pad array on PCB. Therefore, we designed a 32-channel flexible electrode. The number of channels matches the PCB pad array. 32 electrodes are sequentially bonding on PCB to realize the connection of 1024 channel electrodes. During this process, the pad part at the back end of the electrode allows partially overlap. The electrode and the PCB can achieve reliable electrical connection due to the flexibility and toughness of the electrode.

The electrode site is 20 × 20μm, the site spacing is 50μm, and the line width and line spacing are 6μm. The front-end width of the flexible electrode is 330μm. It is slightly smaller than the column spacing of the PCB pads, which can ensure the front ends of the electrodes arranged in sequence without overlapping. The back-end pads of the electrodes are arranged on one side, with a size of 160×200μm. Two holes for gold ball bonding are fabricated on each pad, with a diameter of 60μm. The width of the rear end of the electrode is 620μm, equivalent to the width of the electrode pad plus the electrode wiring.

### 2.5 Fabrication of flexible neural electrode

The preparation process of flexible neural electrodes is as follows: (1) Spin-coating the bottom polyimide (PI) layer and drying; (2) Spin-coating negative photoresist (AR-N4340) and patterning by photolithography; (3) Evaporating Cr/Au/Cr metal layer and removing photoresist to form electrode layer; (4) Spin-coating the top PI layer and drying; (5) Spin-coating positive photoresist (AZ4620) and patterning by photolithography; (6) Exposing electrode sites and pads by reactive ion etching; (7) Repeating steps 5 and 6 to etch out the electrode profile; (8) Corroding the top metal Cr and releasing the electrode.

### 2.6 The interconnection between flexible electrodes and the high-density PCB

Gold ball bonding is used to connect flexible electrodes to the PCB (Fig.2B). Align the pads of the first flexible microfilament with the first row of bonding pads of the PCB and use deionized water for temporary fixation. Gold balls are bonded to the bonding pad through holes reserved in the electrode pads. The diameter of the gold ball is approximately 80μm, which is larger than the hole in the electrode pad, thus allowing for a robust and reliable electrical connection. After bonding the first microfilament, bond the second microfilament with the same procedure. To produce a 1024-channel electrode, a total of 32 flexible microfilaments need to be bonded in sequence (Fig. 4B).

**Fig.4.**
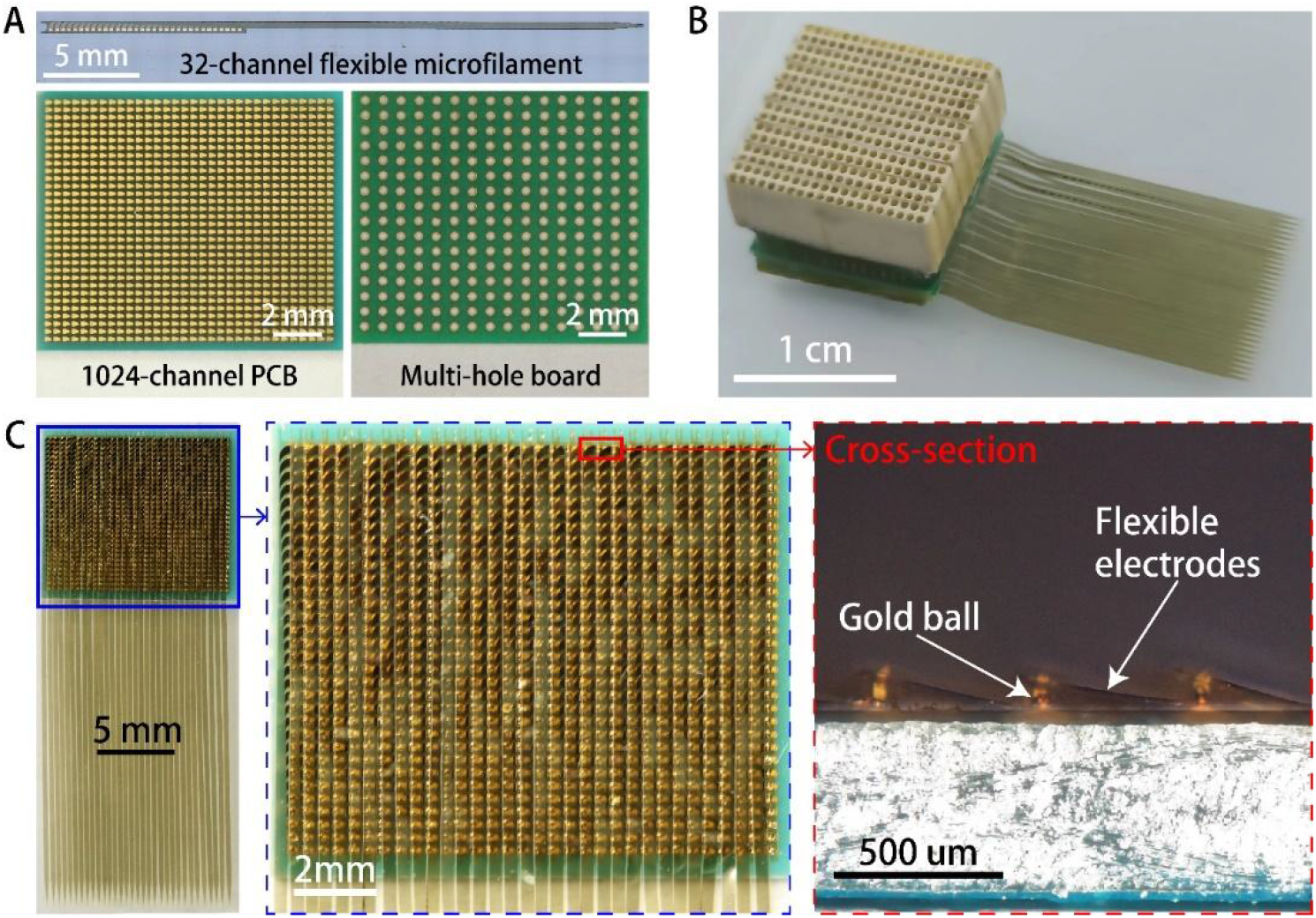
1024-channel flexible neural microelectrodes. (A) 1024-channel PCB and multi-hole boards. (B) 1024-channel flexible neural microelectrodes fan-out. (C) Flexible neural microelectrodes bonding on 1024-channel PCB.

### 2.7 The interconnection between connectors and the high-density PCB

To reliably connect the 256-channel connector module to the 1024-channel PCB, a multi-hole board, like a screen printing plate, was designed to facilitate the process. There are 256 holes on the board, and their spacing and arrangement are matched to the pins of the connector array. The holes are designed to be circular in the first selective fan-out method and are 350um in diameter, slightly larger than the pad on the high-density PCB and the pin of the Omnetics connector. In the second selective fan-out method, the hollow out shape are designed to produce a gating-line on the PCB. The welding process is as follows (Fig.3C): (1) Aligning the multi-hole board with the PCB to expose the 256 pad areas on it; (2) Applying ultraviolet glue to the edges to fix the PCB and the multi-hole board; (3) Coating the solder evenly on the multi-hole board until it fills each hole; (4) Aligning the 256-channel connector module with the multi-hole board, ensuring that each pin is inserted into a hole and submerged by the solder; (5) Placing the aligned PCB and connector modules on a hot plate to evaporate the solvent from the solder (150°C for 10 min); (6) Lowering the temperature to cure the solder and finish the welding.

### 2.8 Electrochemical characterization test of the high-density fan-out method

To characterize the reliability of the proposed high-density fan-out method, the yields of bonding only the electrodes to the PCB and both the electrodes and the connectors to the PCB were tested respectively. Electrochemical impedance (EIS) measured in the phosphate-buffered saline (PBS) solution was used to evaluate the conductivity. The two-electrode system is adopted in electrochemical measurements, with a platinum electrode as the counter and reference electrode. The EIS was measured by applying a sinusoidal AC voltage of 5mV, and the frequency range was 100-10000Hz. The electrode impedance value corresponding to 1KHz is used to evaluate the conduction of the electrode.

## 3 Result

### 3.1 High-density flexible neural electrode packaging

Based on the vertical bonding process and the flexibility of the electrodes, 1024 electrodes on 32 flexible microfilaments are successfully bonded to a PCB in an area of 11.8×10mm^2^, as shown in Fig.4. Almost all areas on PCB are used for interconnection because the whole PCB is spread with pads, and the interconnection density of the 1024 channels can reach 8.8 channels/mm^2^, which exceeds the 3072-channel flexible electrode reported by musk [22] in 2019 (its interconnection density is about 7.2 channels/mm^2^). The front and back ends of the entire electrode array are kept in the same size, which suggests its scalability in the number of channels, and the overall thickness dose not increase too much, due to the ultra-thin flexible electrodes (Fig. 4C).

Based on the proposed selective fan-out method, we have designed the corresponding multi-hole boards (Fig. 4A), and successfully welded 256 from the 1024 channels to the Omnetsics connector module. The bonding and welding pads of the high-density PCB are conducted through the vias, which eliminates the area occupied by the traces on the PCB. And the arrangement of the welding pads of the PCB enables the connector to be tightly and vertically packaged. The volume of the entire back-end packaging is only 12×11×8mm^3^, with the connector module occupied most volume (Fig. 4B). With higher density connectors or chips, the overall package size can be further reduced.

### 3.2 Effective connection rate of high-density packaging method

After bonding the 1024 electrodes to the high-density PCB, the electrochemical workstation is attached to the back of the PCB through a modified multi-probe platform. The impedance and phase spectra of the tested flexible electrodes demonstrate a typical gold-interfaced neural microelectrode (Fig.5 A and B). The effective connection rate of the 1024 channels close to 100% (Fig.5C). And the impedance values of these electrodes showed a normal distribution with a mean of 1.16MΩ (Fig.5D). In addition, based on the second fan-out method, another 256 and 512 channels of 1024 electrodes are selectively bonded to the high-density PCB. The effective connection rate reached both 100%.

**Fig.5.**
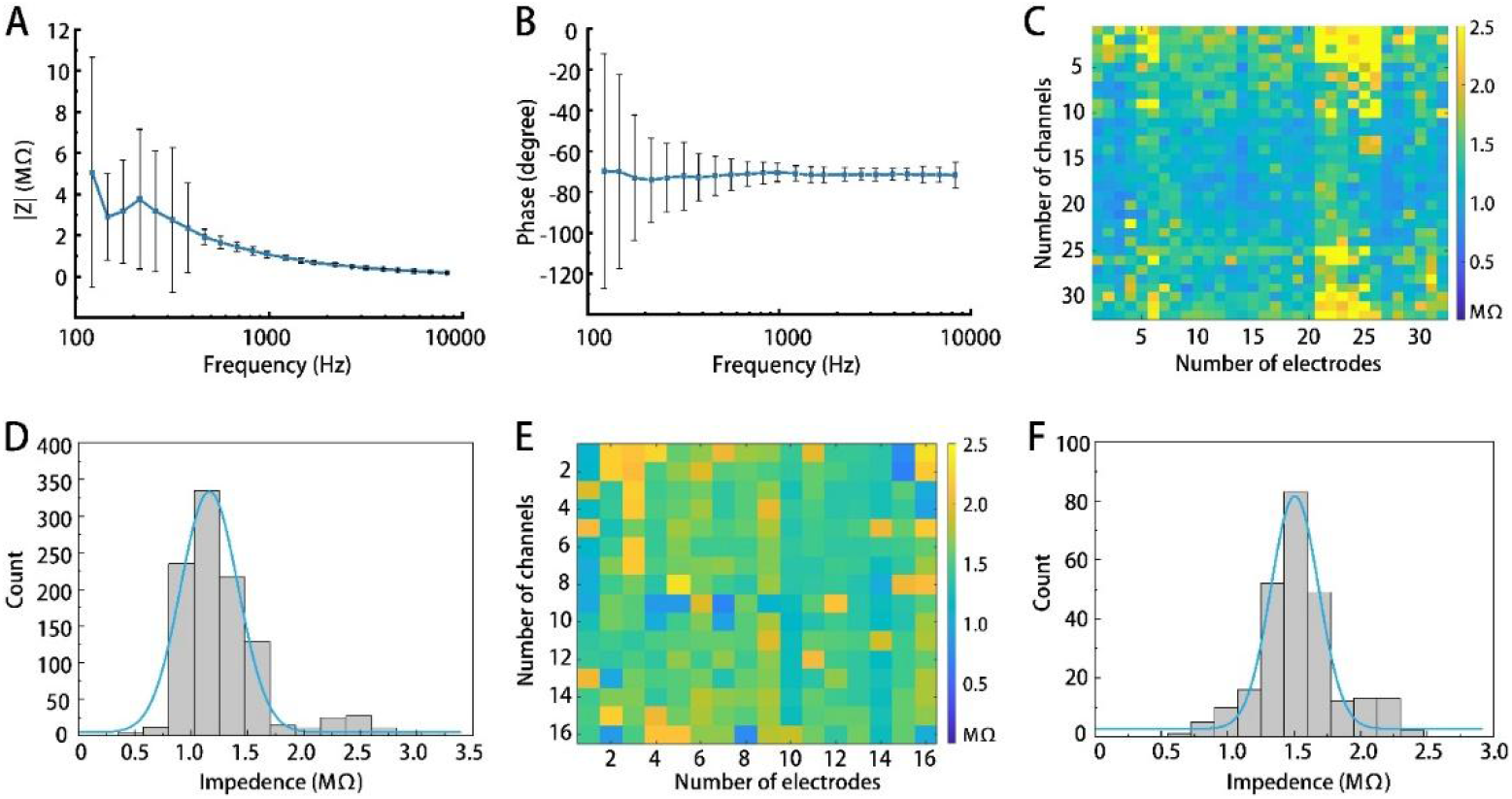
Flexible neural microelectrodes characterization. (A and B) Impedance and phase spectra of the electrode. (C) Impedance of 1024 channels electrodes at 1KHz. (D) Frequency diagram of 1024 channels electrode impedance distribution. (E) Impedance of fan out electrodes at 1KHz. (F) Frequency diagram of fan out electrode impedance distribution.

After welding the 256-channel connector module to the welding pad of the PCB through a multi-hole board, the 256 fanned-out channels are still well connected (Fig.5E). The impedance values also showed a normal distribution with a mean of 1.51MΩ (Fig.5F). which demonstrates the reliability of this proposed high-density fan-out method.

## 4 Discussion

High-throughput flexible neural electrodes can achieve high-density, high-quality neural electrical signal recordings in multiple brain regions with less damage to nerve tissue. However, the back end of the high-throughput electrode packaging is bulky, which seriously affects the recording of electrical signals after implantation. Thus, we propose an ultra-high-density fan-out strategy for high-throughput flexible neural electrodes. We validated this high-density fan-out strategy based on commercially available Omnetsics connectors. 1024 electrodes were successfully connected to the external PCB in an area of 11.8×10mm^2^. And after fanning out 256 of these channels, the entire package is around 1cm^3^ in size. However, these are not the extremes of this fan-out strategy as it has great adaptability and scalability. The number of electrode channels can be easily scaled to much more than 1024 because the size of the PCB increases linearly with the number of electrode channels. The fan-out density can be further increased when there are more compact connectors available. Moreover, if a custom amplifier chip is available, the number of fan-out channels can be greatly increased and the footprint can be further reduced, which indicates great potential of this strategy for application in high-density fan-out of high-throughput flexible electrodes.

## Acknowledgements

The project was supported by the National Key Technologies Research and Development Program of China (2017YFA0701100, 2022YFF1202303); Shanghai Municipal Science and Technology Major Project (2021SHZDZX); the National Natural Science Foundation of China (62071447, 61971049, 22278037); the Strategic Priority Research Program of the Chinese Academy of Sciences Pilot Project (XDB32030102, XDB32040203, and XDA16021305); the Key Scientific Research Project of Beijing Municipal Commission of Education (KZ202010015024, KZ201910015016, KZ202110015019), the Research and Development Program of Beijing Institute of Graphic Communication (Ec202006, Ec202004); Discipline construction of material science and engineering(21090122014).

